# Nanoscale Dynamics of Cellulase TrCel7A Digesting Cellulose

**DOI:** 10.1101/2021.02.18.431891

**Authors:** Zachary K. Haviland, Daguan Nong, Kate L. Vasquez Kuntz, Thomas J. Starr, Dengbo Ma, Ming Tien, Charles T. Anderson, William O. Hancock

## Abstract

Understanding how cellulases catalyze the digestion of lignocellulose is a major goal of bioenergy research. Cel7A from *Trichoderma reesei* is a model exoglucanase that degrades cellulose strands from their reducing ends by processively cleaving individual cellobiose units. Despite being one of the most studied cellulases, the binding and hydrolysis mechanisms of Cel7A are still debated. We used single-molecule tracking to analyze the dynamics of 11,116 quantum dot-labeled *Tr*Cel7A binding to and moving processively along immobilized *Gluconoacetobacter* cellulose. Enzyme molecules were localized with a spatial precision of a few nanometers and followed for hundreds of seconds. Most enzymes bound into a static state and dissociated without detectable movement. Processive enzymes moved an average distance of 39 nm at an average speed of 3.2 nm/s. Static binding episodes preceding and following processive runs were of similar duration to static binding events that lacked any processive movement. Transient jumps of >20 nm were observed, but no diffusive behavior indicative of a diffusive search of the enzyme for a free cellulose strand end was observed. These data were integrated into a three-state model in which *Tr*Cel7A molecules can bind from solution into either a static or a processive state, and can reversibly switch between static and processive states before dissociating. From these results, we conclude that the rate-limiting step for cellulose degradation by Cel7A is the transition out of the static state either by dissociation from the cellulose surface or initiation of a processive run.

## Introduction

Cellulose, which is composed of β-1,4-linked glucan chains hydrogen bonded together into cable-like microfibrils, is the most abundant biopolymer on Earth (3), and is a major source of renewable energy and biomaterials. It can be enzymatically degraded by cellulases that release cellobiose, which is subsequently split into two glucose molecules that can be fermented into biofuels (4). However, the partially crystalline structure of cellulose and its interactions with other components of plant cell walls, such as lignin, make it resistant to enzymatic degradation (5). Optimizing cellulase-dependent degradation of lignocellulosic feedstocks has the potential to improve the cost effectiveness of biofuels; however, this requires a more complete understanding of the mechanisms of cellulose degradation by cellulases. The exoglucanase Cel7A from *Trichoderma reesei* (teleomorph *Hypocrea jecorina*) is a model cellulase enzyme that degrades the glucan chains of cellulose from their reducing ends. Its mechanism of action has been investigated using traditional biochemical methods (6–9), AFM (10,11), single-molecule imaging (12–14), and optical tweezers (15), and these results have contributed to quantitative kinetic models of Cel7A activity (1,2,16,17). Cel7A moves processively at a few nm/s, can operate against an external load of 20 pN, and appears to get stuck in “traffic jams” at high enzyme concentrations (11,15). However, some of these results and the ensuing models are contradictory, leaving uncertainty about how Cel7A in fact operates.

The dominant model used to describe the mechanism of Cel7A involves *i)* adsorption to cellulose, *ii)* complexation with a glucan chain and *iii)* arresting in a blocked state (Fig. 1) (1,2). Initial adsorption from solution to the surface of a crystalline cellulose microfibril may involve the cellulose binding module (CBM) and/or the catalytic domain of Cel7A, and may be static or diffusive (1,2). Complexation involves a glucan chain entering the “tunnel” of the enzyme where the active site lies, and results in processive catalytic hydrolysis of the chain (18–20). From this processive active state, the enzyme may decomplex and return to its initial (inactive) state where it releases the glucan chain and unbinds into solution. Alternatively, it may enter an arrested or “blocked” state. A number of mechanisms can be hypothesized to cause a blocked state: *i)* encountering a molecular “doorstop” on the polymer (e.g., a cross-strand of cellulose or a molecule of hemicellulose or lignin*); ii)* encountering another enzyme, or *iii)* the enzyme reaching the end of a chain (1,2,11). From this blocked state, the enzyme may dissociate from the microfibril surface, or it may have to return to an adsorbed, ucomplexed state before either dissociating from the surface or beginning another processive run.

**Figure 1:**
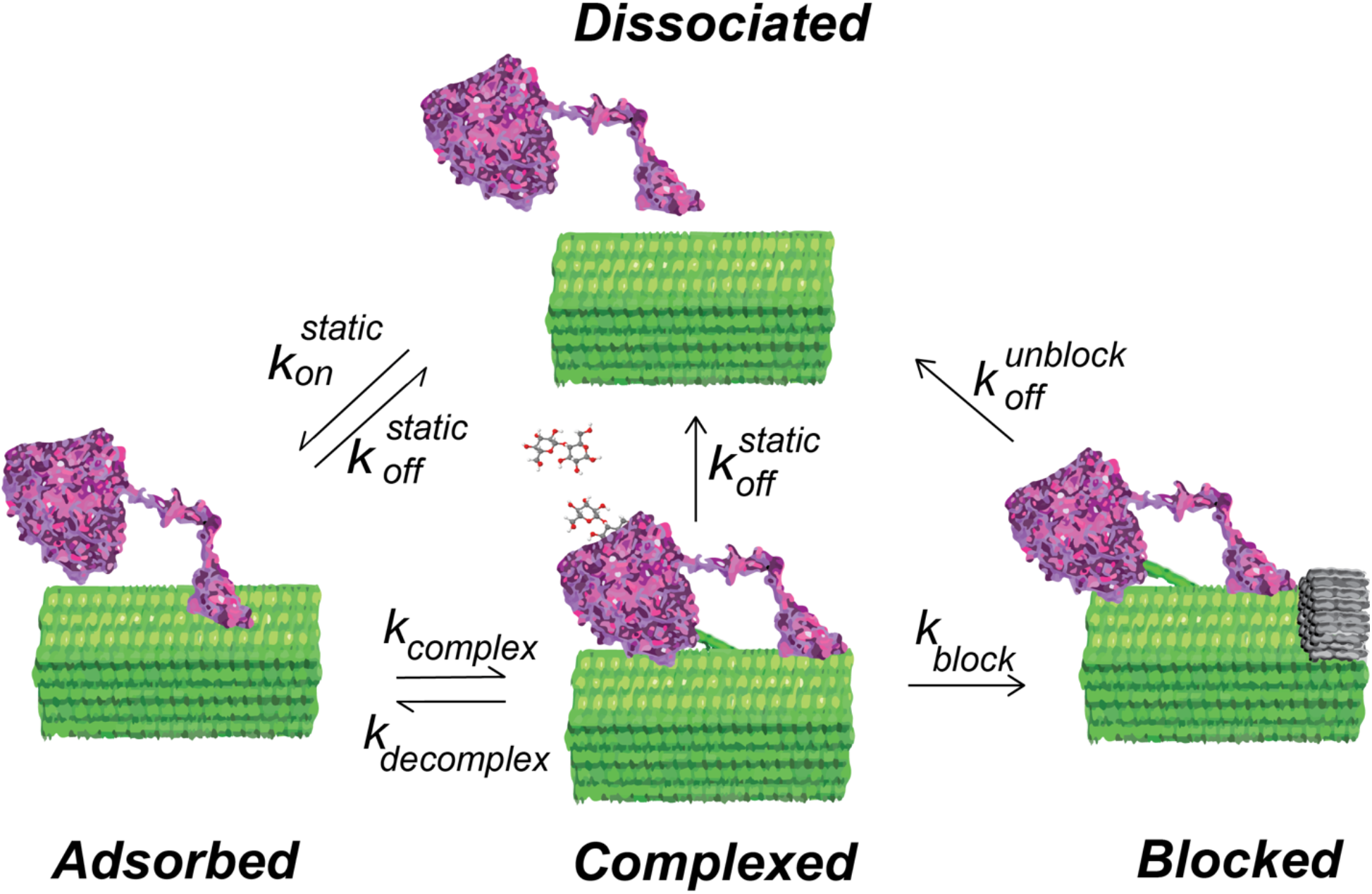
Model for the hydrolytic cycle of Cel7A enzymes on a cellulose substrate (1,2). Enzymes initially adsorb to the cellulose surface and then “complex” with a free reducing end of a glucan chain and enter a processive degradation state. Processive movement along the cellulose is terminated by the enzyme unbinding from the cellulose or entering a “blocked” state.

Despite considerable experimental and theoretical work to date, there are a number of unresolved questions regarding the mechanism of Cel7A: 1) What is the nature of the adsorbed state – is it diffusive or static? 2) What are the dominant modes of initial binding from solution and unbinding back into solution? 3) What limits processivity? One reason these questions are still unresolved is that different techniques are optimized for measuring different aspects of Cel7A molecular function: optical tweezers can detect the steps of individual cellobiose advances and impose a load; high-speed AFM can characterize moving enzymes and roadblocks; and steady-state kinetic assays provide values for the overall turnover number (k_cat_) (8,11,15). Single-molecule fluorescence investigations have provided estimates of binding times, but the limited spatial resolution achieved to date using fluorescence has constrained the degree of insight into the processive hydrolysis dynamics of Cel7A. In addition, the bleaching inherent in organic fluorophores has limited the temporal resolution and duration of fluorescence tracking experiments (12–14,21).

Here, we used Interference Reflection Microscopy (22,23) to image unlabeled cellulose, and tagged Cel7A with quantum dots that provide excellent signal/noise and minimal bleaching for visualization by Total Internal Reflection Fluorescence Microscopy. Using a custom-build SCATTIRSTORM microscope, we tracked 11,116 Cel7A molecules interacting with bacterial cellulose immobilized on glass coverslips. We found that most Cel7A molecules bind for tens of seconds and dissociate without moving measurably along the cellulose. A subset of the molecules bind and alternate between static and processively moving states. We combined our measurements into a three-state model whereby Cel7A can bind into or unbind from either a static state or a processive state, respectively. The enzymes occasionally make rapid jumps on the substrate, but we did not observe any evidence that the enzymes diffuse along the substrate while searching for a free reducing end to digest. These results provide new constraints for modeling the enzymatic mechanism of Cel7A.

## Results

### Isolation of bacterial cellulose and Cel7A preparation

A range of cellulose preparations were investigated, including acid-treated *Cladophora* cellulose and commercial AviCell from wood pulp. Based on the ability to adhere to glass slides, ease of handling, and behavior in microscopy studies, we found that bacterial cellulose from *Gluconoacetobacter hansenii* was an ideal substrate. Previous studies have shown that this bacterial preparation exhibits higher crystallinity than plant cellulose(24–26). Cellulose collected from *G. hansenii* cultures was purified by base-treatment, sonicated and then treated with a microfluidizer as described in Materials and Methods. The average concentration of total sugars in the cellulose sample was determined to be 4.58 mM. The degree of polymerization was determined to be 292 by dividing the total sugar concentration by the concentration of reducing ends, determined using a bicinchoninic acid colorimetric assay (27). Purified *Trichoderma reesei* Cel7A (Sigma) was biotinyated using biotin-NHS (Thermo-Scientific), and incubated with streptavidin-coated Quantum dots (Qdots; 525 nm emission; Thermo-Scientific).

### Imaging Cel7A on immobilized cellulose

To investigate the binding characteristics of Cel7A, we adsorbed cellulose to plasma-treated glass coverslips, assembled flow cells using double-sided tape, and imaged Qdot-labeled Cel7A interacting with the surface-immobilized cellulose. Following cellulose adsorption, surfaces were blocked by flowing 1 mg/mL bovine serum albumin (BSA) into the flow cell for 5 min, followed by an enzyme solution containing 2 or 10 nM Cel7A combined with 0.5 nM Qdot in 50 mM sodium acetate buffer plus 5 mM dithiothreitol, pH 5.0. The Qdot-labeled Cel7A was imaged by Total Internal Reflection Fluorescence Microscopy (TIRFM) using a 488 nm laser (Fig. 2A) on a custom-build microscope (28). Tetraspeck beads (Thermo Scientific) (brighter objects in Fig. 2A) were imaged simultaneously as fiduciary markers to compensate for stage drift (29). Immobilized cellulose was imaged by Interference Reflectance Microscopy (IRM) (22,23,28) (Fig. 2B). In time-lapse experiments, Qdot-labeled Cel7A was observed reversibly binding to the immobilized cellulose. As seen in the merged image in Fig. 2C, Cel7A bound preferentially to the immobilized cellulose and non-specific binding to the BSA-blocked glass was negligible.

**Figure 2:**
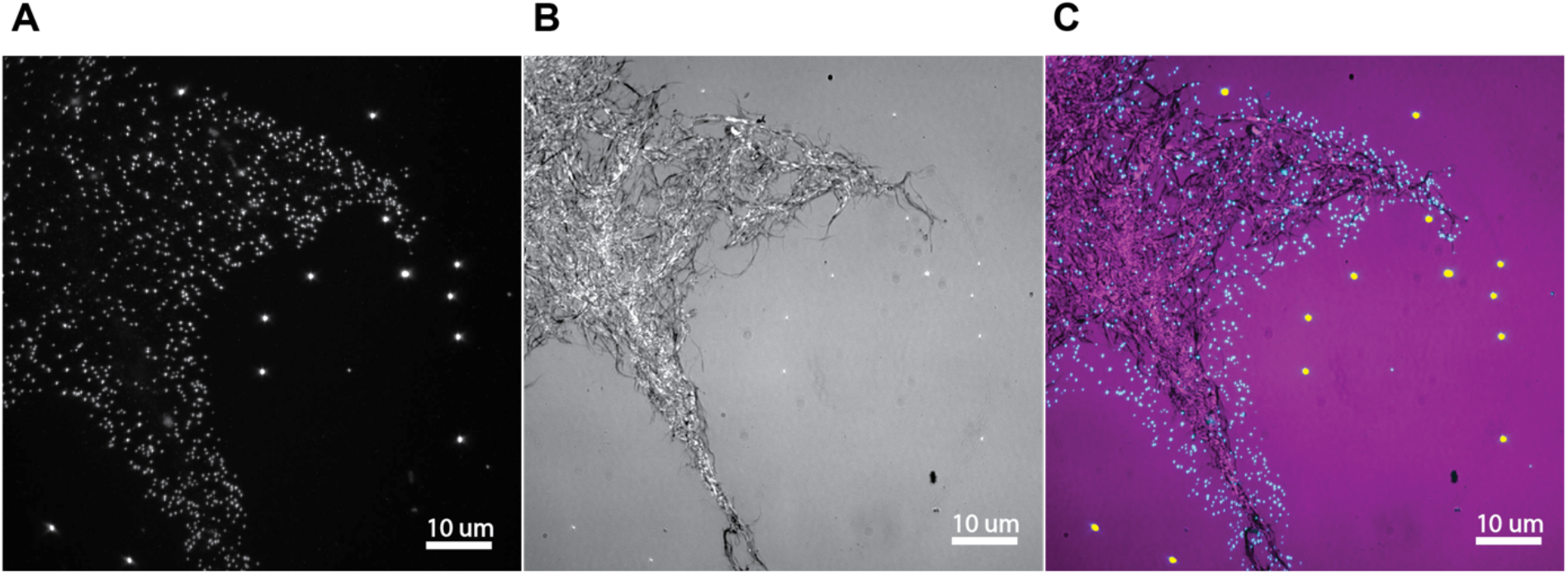
Qdot-labeled Cel7A enzymes reversibly interact with surface-immobilized cellulose. A: Total Internal Reflection Fluorescence of Qdots and Tetraspeck beads (bright objects away from the cellulose). B: Interference Reflection Microscopy image of bacterial cellulose adsorbed to glass coverslip. C: Overlay of fluorescence and IRM images (Qdots are colored cyan while Tetraspeck beads are colored yellow). See also Supplemental Movie 1.

To analyze the dynamics of Cel7A during cellulose degradation, 1000-sec movies at 1 frame/sec were recorded, and Qdot positions were localized by fitting a 2D Gaussian point-spread function using FIESTA software (30). Changes in the positions of the Tetraspeck particles were subtracted to compensate for stage drift in the x-y plane, and drift in the z-direction was minimized by an active feedback system on the microscope. A custom analysis program written in Matlab was used to analyze particle trajectories (28). The tracking precision of the system was determined by immobilizing a Qdot525 on a coverslip, moving the piezoelectric stage in a stepwise pattern, and determining particle positions by the tracking software. The particle positions accurately reflected the step displacements, and the standard deviation of position at each step was 1.5 nm (28). Using this system, we analyzed the binding dynamics and movement of 11,116 Cel7A molecules across two independent experiments.

### Cel7A dynamics on immobilized cellulose

When imaging Cel7A on immobilized cellulose, we identified three types of behaviors: static binding, processive movement, and transient jumps (Fig. 3). The majority of the enzymes (89.9%) that bound to the cellulose substrate remained static throughout their binding duration before eventually dissociating back into solution. These “static” molecules were defined as moving less than 10 nm from their original binding site, and the minimum duration to be counted as an event was 10 s. The durations of the static Cel7A binding events were roughly exponentially distributed (Fig. 4) with a mean binding duration of 89.0 s (95% confidence interval 85.3 - 92.7 s, N = 4,136 molecules). Notably, the enzymes were not observed diffusing along the surface of the cellulose substrate as might be expected if they were bound only through the cellulose binding module (31).

**Figure 3:**
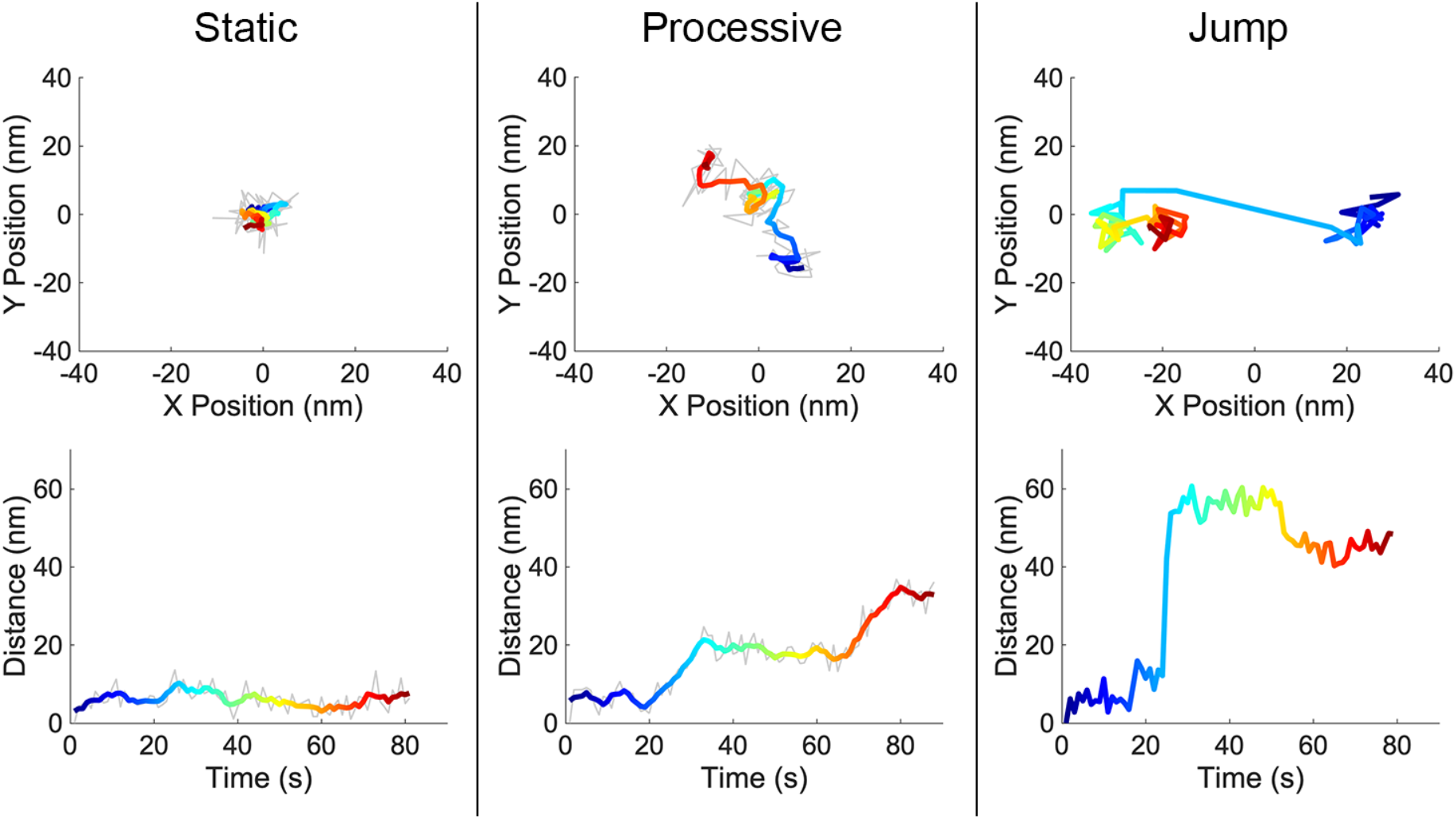
Single-molecule trajectories of individual Cel7A proteins on immobilized cellulose. Qdot-labeled enzymes were imaged at 1 frame/sec, and position was localized by fitting a Gaussian point-spread function. Top row: x-y positions over time; bottom row: distance from origin versus time for the same proteins. Static enzymes (left) are those that move <10 nm over their entire bound duration. Processive enzymes (middle) are those that move >10 nm over a duration of >5 s. Jumps (right) are displacements of >10 nm over two frames, which are interpreted as dissociation and diffusion through solution, followed by rebinding. The colored lines in the static and processive plots are a 5-point boxcar average of the raw data, with time color-coded from blue to red. The colored lines in the jump tracks show the actual position of the enzyme to emphasize their rapid displacement. See Figure S1 for further example traces.

**Figure 4:**
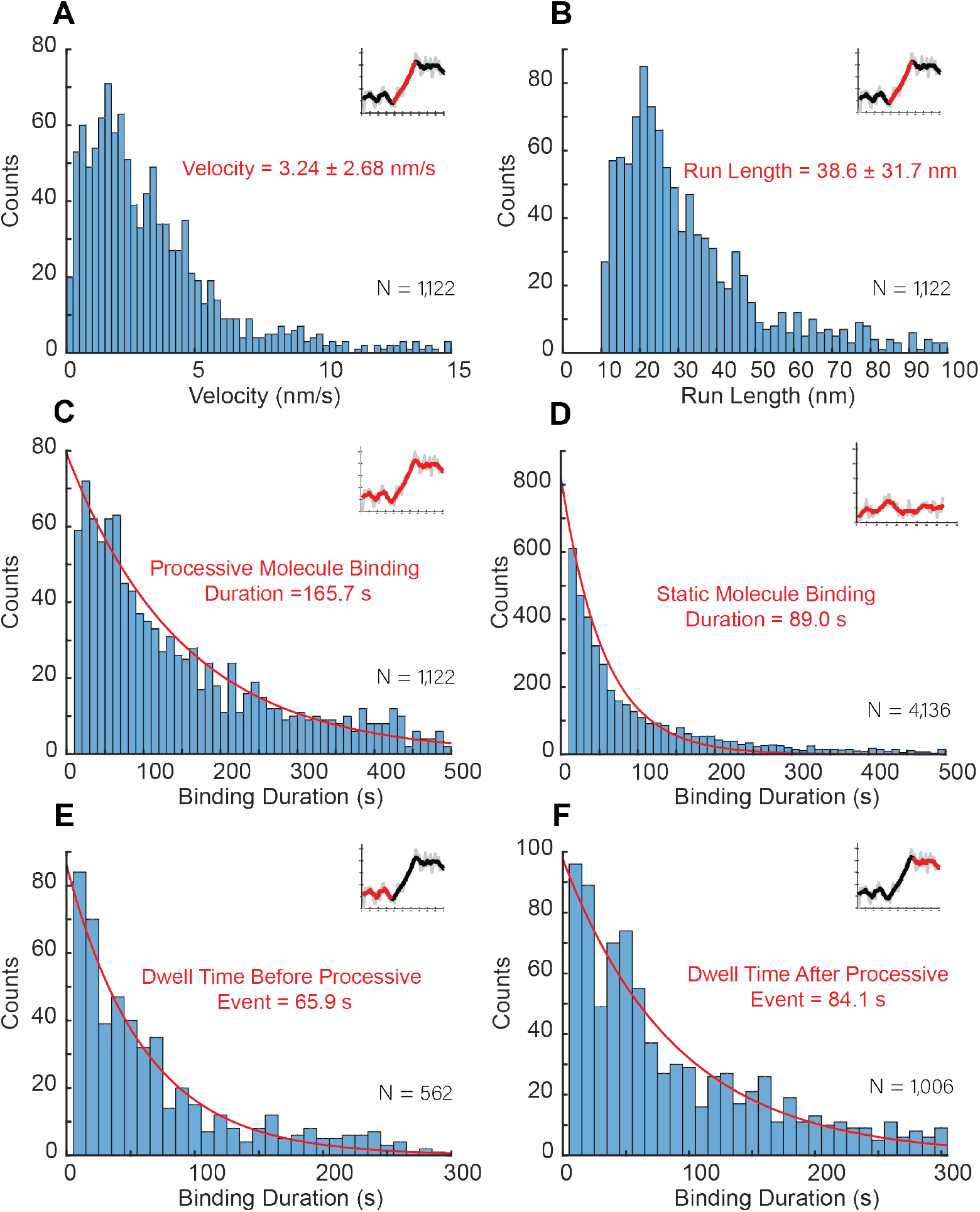
Binding and moving characteristics of sub-populations of Cel7A enzymes. A: Velocity of processive motility segments. B: Run length of processive motility segments; segments below 10 nm were not counted. C: Binding duration of processive molecules, which includes switching between processive and static segments. D: Binding duration of static molecules. E: For processive molecules, dwell times of static segments that preceded processive segments. F: For processive molecules, dwell times of static segments that followed processive segments. Velocity and run length were calculated as mean +/− SD, and binding durations were fit to a first-order exponential function, as described in text. See also Fig. S2 for single plot of all static durations and Fig. S4 for binding duration of all static events.

Processive movement, defined as an overall displacement of 10 nm or more over a duration of at least 5 s, was seen in 10.1% of the enzymes. In addition to the observed displacements, most processive enzymes also had at least one static segment, which was defined as a period (> 5 s) during which displacement was less than 10 nm (Fig. 3 and S1). The mean velocity of the processive segments was 3.24 +/− 2.68 nm/s (mean +/− SD, N = 1,122 segments), in reasonable agreement with previous work (10–12,15). The mean duration of these processive segments was 20.3 +/− 38.9 s (mean +/− SD, N = 1,122 segments), and the mean displacement during these processive segments was 38.6 +/− 31.7 nm (mean +/− SD, N = 1,122 segments) (Fig. 4 and Table 1). The mean binding duration of these processive enzymes was 165.7 s (95% confidence interval 151.7 - 189.0 s, N = 1,122 molecules), which is longer than the binding duration of the static enzymes, and is consistent with their containing both static and processive segments. To test whether the static segments of processive molecules were analogous to those of entirely static molecules, we measured the durations of these segments. Static segments preceding processive runs averaged 65.9 s (95% confidence interval 60.0 - 61.7 s, N = 562 segments), and static segments following processive runs averaged 84.1 s (95% confidence interval 78.4 - 90.2 s, N = 1,006 segments) (Fig. 4 and Table 1). These durations were within a factor of two of one another, consistent with the static segments separating processive runs being mechanistically identical to the purely static binding events (see Figure S2 for plot of superimposed static distributions).

**Table 1:** Complete set of measured parameters from single-molecule experiments.

| Parameter | Value | Units |
| --- | --- | --- |
| All Molecules: |  |  |
| Total number of molecules | 11,116 |  |
| Percent entirely static | 89.9% |  |
| Binding duration of entirely static | 89.0 (85.3 – 92.7)* | s |
| Percent with processive segments | 10.1% |  |
| Binding duration of processive molecules | 165.7 (151.7 – 189.0)* | s |
| Processive Molecules: |  |  |
| Percent bound into processive state | 56% |  |
| Percent bound into static state | 44% |  |
| Number of processive segments | 1,618 |  |
| Duration of processive segments | 20.3 ± 38.9 | s |
| Velocity | 3.24 ± 2.68 | nm/s |
| Run length | 38.6 ± 31.7 | nm |
| Percent of processive segments that end in unbinding | 23% |  |
| Duration of processive segments that end in unbinding | 18.6 ± 17.4 | s |
| Percent of processive segments that end in static event | 77% |  |
| Duration of processive segments that end in static event | 21.4 ± 25.2 | s |
| Static Segment of Processive Molecules: |  |  |
| Number of static segments | 1,820 |  |
| Dwell time of static segments | 85.9 (79.8 – 92.1)* | s |
| Percent of static segments that end in unbinding | 63% |  |
| Duration of static segments that end in unbinding | 84.1 (78.4 – 90.2)* | s |
| Percent of static segments that end in processive event | 37% |  |
| Duration of static segments that end in processive event | 65.9 (60.0 – 71.7)* | s |
| Transient Jumps: |  |  |
| Distance | 48.6 ± 42.5 | nm |
| Binding duration of molecules that jump | 186.3 (163.9– 213.5)* | s |
| Percent of molecules with jumps | 7.3% |  |

\* Values calculated using 95% confidence interval of single exponential fit
± Values calculated using mean and standard deviation

The third behavior observed was “jumps”, which were defined as a displacement of more than 10 nm within two frames (< 2 s) (Fig. 3). These jumps were observed in 7.3% of molecules analyzed and were observed for both static and processive molecules. To confirm that the jumps were not rapid processive segments, movies were taken at a frame rate of 10 frames/sec. These rapid displacements were again observed within two frames (200 ms) (Fig. S3). A plausible mechanism for these jumps is the molecules dissociating from the cellulose, briefly diffusing in solution, and then reassociating within 100 nm of their original binding site. The search space was limited to 100 nm to increase the probability that the same molecule was being observed dissociating and reassociating, and it is likely there were many jumping events with larger displacements that escaped detection. A subset of these jumps (42.5%) occurred immediately upon the enzyme landing on the cellulose substrate, which might represent a “search” behavior for a free glucan chain analogous to a diffusive search, but involving full unbinding rather than diffusion along the cellulose surface while bound. However, these rapid jumps were rare events that involved transient interactions with the surface rather than diffusion along the surface. Based on these three distinct behaviors, various dwell times and run lengths were calculated to determine any differences between the subpopulations.

### A three-state model of Cel7A

To better understand the relationship between static and processive states during cellulose degradation by Cel7A, we constructed a three-state model to describe our kinetic data (Fig. 5). In the model, enzymes can be freely diffusing in solution, statically bound to the substrate, or processively degrading the cellulose substrate. To define the rate constants in our model, we used the durations of the static binding events and processive segments, the probability of switching between static and processive state, and the rate of unbinding from cellulose from Table 1. As shown in Fig. 4, the static binding durations and the durations of the processive runs were all exponentially distributed. This property is consistent with the exits from these states being defined by first-order rate constants (32). The first-order transitions out of these states can thus be calculated by taking the inverse of the binding duration time constant. Using the weighted average of the static states, the first-order transition rate out of the static state is 0.0113 s^−1^. Similarly, based on the 20.3 s mean duration of the processive segments (Table 1), the first-order transition rate out of the processive state is 0.0493 s^−1^. Note that both the static and processive states have two possible exit routes – dissociation or transition to the other bound state (Fig. 5). Accordingly, the transition rates out of these states represent the sum of the two rate constants. For instance, the 0.0493 s^−1^ transition rate out of the processive state is the sum of k_off_ and k_static_ in Fig. 5. The relative values of these two exit routes from the processive state can be calculated as follows: from Table 1, 23% of the processive segments terminated due to the enzyme unbinding from the cellulose and 77% of the processive segments terminated due to the enzyme entering the static state. Thus, k_off_processive_ is 0.23 * 0.0493 s^−1^ = 0.0113 and k_static_ is 0.77 * 0.0493 s^−1^ = 0.0379. A similar calculation can be made for the static states, as follows: 89.9% of the total binding events consisted static binding and dissociation, and among all of the processive enzymes there were 1,820 static segments, of which 63% terminated by the enzyme dissociating from the substrate and 37% terminated due to the enzyme entering the processive state. This comes out to an off-rate from the static state, k_off_static_ of 0.0106 s^−1^ and a transition rate into the processive state, k_processive_ of 0.00064 s^−1^.

**Figure 5:**
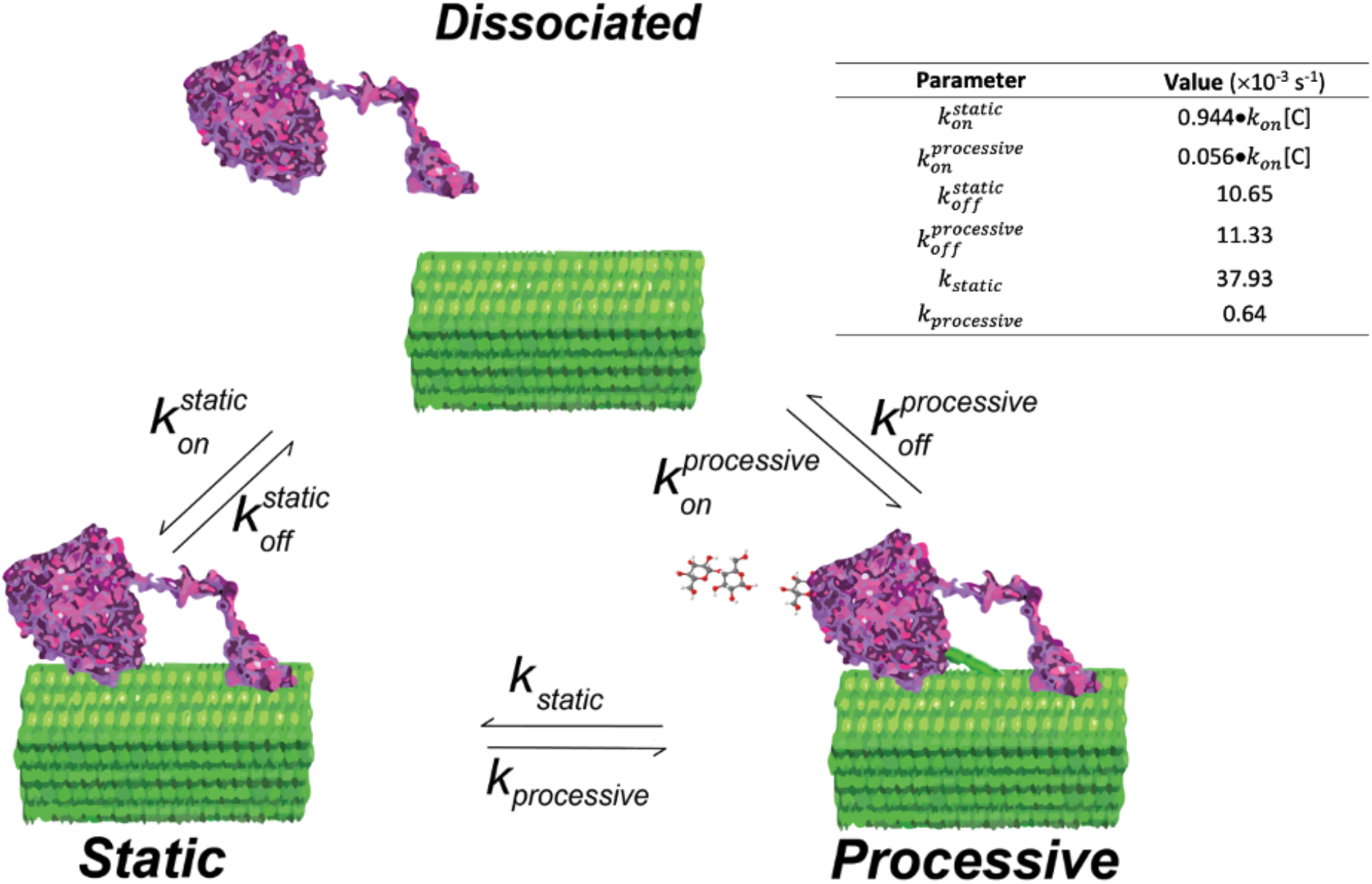
Three state model that accounts for the single-molecule observations. Cel7A in solution can bind to cellulose in a static state or into a processive state. While bound to the cellulose, enzymes can switch between the static and processive states, and enzymes can dissociate from either the static or processive state. Kinetic parameters were calculated from dwell times in the static and processive states, and probability of switching states versus dissociating. The bimolecular on-rate, kon, cannot be determined from these measurements, but the relative on-rate into the static and processive states are calculated by the probability of the initial state being static or processive.

The above analysis provides four of the six rate constants in the model, leaving the two on-rates undetermined. Because of uncertainties for true cellulose concentration in the flow cell and the fact that cellulose is a heterogeneous and insoluble substrate, we are unable to measure the true bimolecular on-rate in our system. However, we are able to determine the relative probability of landing into the processive or static states, as follows. 89.9% of binding events were purely static, and of the processive molecules, 44% started with a static segment before starting to move whereas 56% began moving immediately upon landing on the cellulose (Table 1). Thus, 94.4% of binding events entered a static state and the on-rate into the static state is ~20-fold higher than the on-rate into the processive state. This observation shows the static state accounts for most of the binding events on the substrate from enzymes in solution and emphasizes how often the enzymes are in the static state.

## Discussion

### What is the mechanism of cellulose digestion by Cel7?

By tracking Cel7A with nm precision, we find that enzyme molecules bind reversibly to immobilized crystalline cellulose and transition between static and processive states with characteristic times of tens of seconds. The finding that static states are long-lived is consistent with the existing paradigm that the off-rate of the enzyme from the substrate is rate limiting in the reaction (8,33,34). Existing models generally separate adsorption to crystalline cellulose surface from complexation of the glucan chain entering the tunnel of the enzyme, and they introduce a “blocked” state that terminates the processive run (1,2). However, our results are consistent with a simpler model (Fig. 5) that includes a single static state where the enzyme is bound to the substrate and a single processive state where the enzyme is hydrolyzing the substrate.

Previous single-molecule fluorescence studies have observed reversible binding of Cel7A to cellulose, but with somewhat different kinetics and different interpretations. Shibafuji et al (13) measured a biexponential distribution of binding durations, with characteristic time constants of approximately 1 s and 8 s, and attributed the shorter time to nonproductive binding and the longer to productive hydrolysis. Jung et al (14) measured characteristic binding times of 30 s and 173 s and similarly attributed the longer time to productive binding events. Here, we separated static binding events (characteristic time of 89.0 s) from processive events (characteristic time 165.7 s including both static and processive interludes). In this way, our durations agree more closely with Jung et al (14). However, by recognizing that the processive molecules included static phases with similar durations to the strictly static binding events, we do not classify individual molecules into productive and non-productive events. Instead, we interpret the purely static events as binding events where the enzyme dissociates before beginning a processive run. One corollary of this is that, because enzymes can switch from a static state into a processive state and back, enzymes that move processively will tend to have longer total binding durations because they include a processive segment and one or more static segments. A small number of the enzymes we imaged (0.162% of entire population) moved immediately upon binding and dissociated from this moving state; the vast majority of processive enzyme had at least one static segment.

The processive movement we observe is consistent with hydrolysis-driven movement visualized and inferred by others. Processive Cel7A movement has been most clearly observed by high-speed AFM, where velocities averaged 3 - 7 nm/s in related studies (10,11). An optical trapping study measured a mean velocity of 0.25 nm/s (15); these values bracket our 3.2 nm/s mean velocity, and a wide distribution of velocities is seen across different techniques. A kinetic analysis of cellulose degradation by Cel7A (8) calculated a predicted turnover rate (i.e., production rate of cellobiose) of 4 s^−1^ during processive degradation, agreeing qualitatively with our measured velocity and a cellobiose length of ~1 nm, and predicted a processive burst of 13 cellobiose subunits, which corresponds to about a third of our average run length of 39 nm. Notably, this enzymatic analysis also calculated that at steady-state only ~10% of bound enzymes are productively hydrolyzing cellulose, consistent with our observation that the majority of binding events do not result in processive movement.

### What is the mechanistic interpretation of the static state?

The static state we observe could potentially represent tethering of Cel7A to the cellulose through its cellulose binding module (CBM), or it could represent engagement of the catalytic domain with a cellulose strand without processive hydrolysis. Although we cannot definitively rule out either possibility, three lines of evidence argue for nonproductive engagement of the catalytic domain rather than tethering through the CBM. First, diffusion rates of isolated CBMs on cellulose have been found by FRAP measurements to be ~2 ϗ 10^3^ nm^2^/s (31). In the 89.0 s mean bound duration we measured, a molecule with this diffusion constant would be expected to diffuse laterally across a mean distance of ~ 850 nm (<x^2^> = 4Dt) (35). In contrast, our static particles were restricted to <10 nm displacements during their binding durations. Second, a recent report used fluorescence unquenching of Trp residues in the substrate tunnel to monitor binding of Cel7A to crystalline cellulose and found a consistent single-exponential rise to a steady-state fluorescence with a calculated off-rate of 0.005 s^−1^ (6). The lack of a lag phase in that work suggests that there is no delay between binding of Cel7A to cellulose and threading of a glucan chain into the substrate tunnel, and their measured off-rate corresponds to a 200 s binding duration, which is close to our observed processive binding durations. Third, a number of studies have compared the activity of isolated Cel7A catalytic domains lacking a CBM to the intact enzyme and found that deleting the CBM results in only minor changes in binding affinity and activity (12,15,21). Thus, although we cannot rule out the CBM assisting in binding and processive hydrolysis, we interpret the static states we observed as representing unproductive engagement of the catalytic domain of Cel7 with a cellulose strand.

### What events terminate the processive hydrolysis of cellulose by Cel7A?

We found that during processive movement, Cel7A molecules moved an average of 39 nm, corresponding to ~39 cellobiose molecules being released (Fig. 4B and Table 1). The degree of polymerization of our bacterial cellulose was found to be ~300, which suggests that processive runs are likely not terminated by the enzyme reaching the end of the glucan chain it is degrading. One caveat is that if Cel7A preferentially interacts with short glucan chains, which might be expected to be preferentially found at the exposed surface of the cellulose, then a significant fraction of processive runs could be terminated by end of the chain. Approximately 23% of processive runs ended with the enzyme dissociating from the cellulose (Table 1), likely due to either dissociation from chain termini or chain “dethreading” from the substrate tunnel of the enzyme followed by unbinding of the enzyme from the cellulose surface (33). 77% of processive runs ended with the enzyme stalling on the cellulose; the mechanism underlying these stalls is not clear. The possibility that processive cellulose degradation stalls due to Cel7A encountering other enzymes on the substrate has been investigated in detail using high-speed AFM (11). However, we rule out enzyme “traffic jams” as the source of the pausing behavior in the current work because our study used enzyme concentrations that were three orders of magnitude lower than those in the AFM work. Furthermore, stalling was observed in our work when no other enzymes visible in the vicinity (see low particle density in Fig. 2). Stalling could potentially occur from product (cellobiose) inhibition, but this is unlikely because stalls were observed even in the earliest landing events during an experiment, when cellobiose concentration is very low. Moreover, the K_I_ for cellobiose is estimated at ~0.2 mM, far out of the range of potential product build up for the nM enzyme concentrations used in our experiments(34). Instead, the most likely explanation is that the static episodes result from the enzyme being unable to extract the glucan chain it is hydrolyzing from the crystalline lattice of the cellulose. The fact that static segments following processive runs had similar durations as static segments preceding processive runs and completely static traces suggests that they all represent a similar state of the enzyme. In this way, initial binding of Cel7A and stalling at the ends of processive runs are similar states in that the glucan chain is at least partially engaged in the substrate tunnel, and the engaged chain is still mostly incorporated into the crystalline lattice.

### Do diffusive processes influence cellulose degradation by Cel7A?

The simplest mechanism to envision for the initial encounter of Cel7A with crystalline cellulose is diffusive engagement through the CBM followed by productive engagement of the catalytic domain. However, we did not observe evidence for such a diffusive search process. We cannot rule out search encounters shorter than our 1 s frame acquisition time. Also, we define a static molecule as having a positional standard deviation of < 10 nm, so we cannot rule out that the long static segments we observe involve a very localized diffusional process under this limit. We did observe transient jumps of tens of nm. The predicted diffusion constant of a free enzyme in solution is four orders of magnitude faster than the reported diffusion rate of isolated CBMs on cellulose (35). Thus, if enzymes were rapidly unbinding, diffusing through solution, and rapidly rebinding at rates faster than we are observing, this could be an efficient search mechanism that involves diffusion through solution rather than diffusion along the crystalline substrate.

## Conclusion

We found that *T. reesei* Cel7A degrades bacterial cellulose by alternating between a relatively long-lived static state and a shorter duration processive state. The enzyme can land on the cellulose substrate in either state and can also dissociate from the cellulose in either state. On bacterial cellulose, Cel7A spends the bulk of its time (>95%) in the static state, meaning that the maximal steady-state velocity (k_cat_) is expected to be more than an order of magnitude smaller than the instantaneous degradation rate, in agreement with previous kinetics studies (8). This model is broadly consistent with the idea that dissociation from cellulose is the rate-limiting step of the enzyme (8,33,34). However, in our scheme, Cel7A can land on cellulose in either the processive or the static state, and the enzyme can exit this static state either by dissociation into solution or by transition to processive degradation. Thus, it is exit from the static state that limits the overall efficiency of the enzyme. Accordingly, it is expected that Cel7A will have a higher k_cat_ in bulk assays using substrates in which glucan chains are more easily extracted from the bulk lattice. Conversely, Cel7A may work more slowly on complex substrates such as plant cell walls that include hemicellulose and lignin if these components limit the ability of the enzyme to extract a cellulose strand from the crystalline lattice. Our results point toward new strategies for engineering Cel7A to optimize its hydrolysis of cellulose for the generation of bioenergy and other useful products.

## Materials and Methods

### Cellulose preparation and characterization

We first inoculated Schramm-Hestrin medium (36) with *Gluconacetobacter hansenii* (strain ATCC 23769) and allowed the culture to grow for 5 days at 30 °C with no agitation. The resulting sheet of cellulose was washed five times with 100% ethanol. After filtration, 2% (w/v) NaOH was added to the cellulose and the solution was incubated for 30 min at 80 °C. Next, the solution was centrifuged for 15 min at 2,300 rcf and the supernatant was decanted. To neutralize the pH, the solution was washed once with 0.5 M NaOAc and twice with sterile ddH_2_O. and air dried for 2 days on aluminum foil. After the cellulose was thoroughly dried to a flaky texture, it was peeled off and stored at 4 °C. Dried cellulose was re-suspended in 50 mL ddH_2_O and sonicated with a Sonic Dismembrator (Thermo Fisher, model 100) five times for 30 seconds each at a setting of 9, with 1 min breaks in between. Sonicated cellulose samples were combined and processed through a M-110EH microfluidizer at the Pennsylvania State University CSL Behring Fermentation Facility. The sample was first passed through a 200 μm filter five times at 5000 psi and then passed through a 75 μm filter for 45 min at 7000 psi. The cellulose content was determined by phenol sulfuric acid using a glucose standard (37). The reducing end concentration was determined by the bisynchonic acid method (27). The degree of polymerization was calculated by dividing the total sugar concentration by the reducing end concentration.

### Cel7A preparation and characterization

*T. reesei* Cellobiohydrolase I (Sigma-Aldrich, catalog number: E6412), hereafter referred to as Cel7A, was buffer exchanged into 50 mM NaCl using a PD-10 column (General Electric). Peak fractions, as determined by absorbance at 280 nm, were pooled and 100% glycerol added for a final concentration of 30% (v/v). The final protein concentration (14.8mM) was determined by absorbance, using an extinction coefficient of 74,906 M^−1^cm^−1^. Protein was divided into 200 μl aliquots and stored at −20 °C. After thawing for experiments, enzymes were never refrozen.

Cel7A was biotinylated using EZ-Link NHS-LC-LC-Biotin (Thermo Scientific, catalog number: 21343), which labels the primary amines of exposed lysine residues. Cel7A was buffer exchanged into ddH_2_O, and borate buffer (pH 8.5) was added to make a final concentration of 45 mM NaBO_3_. Biotin dissolved in dried Dimethylformamide (DMF) was added to the Cel7A mixture with a biotin:enzyme ratio of 10:1 and incubated for 6 h in the dark at 21°C. To remove the free biotin, the enzymes were then buffer exchanged into 50 mM sodium acetate buffer using a PD-10 desalting column. The enzyme concentration was calculated using absorbance measurements at 280 nm and the biotin concentration was determined using the Pierce Fluorescence Biotin Quantitation Kit (Thermo Scientific, catalog number: 46610). Biotinylation fraction was determined to be 60%. Biotinylated enzymes were flash frozen using liquid nitrogen and stored at −80 °C.

### Single-molecule Imaging and Analysis

To prepare flow cells, a ~10 ul volume of 0.16 mg/mL of acetobacter cellulose in 50 mM sodium acetate (pH 5.0) was pipetted onto the surface of a plasma-cleaned glass slide. Two strips of double-sided tape were positioned on either side of the cellulose solution and a plasma cleaned glass cover slip was placed on top of the tape to create a flow cell (~45 μL volume). The slide was inverted and placed into an oven at 65°C for 40 min to allow the cellulose solution to dry, leaving the cellulose fibers stuck to the surface of the cover slip. Tetraspeck beads (Thermo Scientific) as fiduciary markers were flowed into the flow cell and incubated for 3 min to allow them to nonspecifically bind to the glass surface. This was followed by three washes of 1 mg/mL bovine serum albumin with three minutes incubation each, to prevent nonspecific binding of cellulase enzymes to the glass surface.

Qdot-labeled Cel7A was prepared by mixing 2-10 nM Cel7A with 0.5 nM Qdot 525 (Thermo Scientific) in 50 mM sodium acetate, pH 5.0, with 5 mM dithiothreitol to prevent photobleaching. Following a 15 min incubation, the solution was injected into the flow cell. Decreasing the enzyme:particle ratio below this led to many fewer landing events, consistent with Qdots binding single enzymes (38). Identical results were obtained using 2 nM and 10 nM enzyme, and so data for the two concentrations were pooled for the final analysis. Single-molecule imaging was carried out through total internal reflection fluorescence microscopy with an excitation laser of 488 nm at 10 mW power to illuminate both the Tetraspeck beads on the surface and the Qdots attached to the enzymes. Cellulose was visualized by interference reflectance microscopy with a 520 nm LED laser. Images were taken at 1 frame/sec and videos consisted of 1000 frames. The imaging area for each frame was 79.2 μm ϗ 79.2 μm with a pixel size of 66.0 nm. A QPD sensor connected to the microscope stage prevented drift in the z-direction to keep the images in constant focus. All videos were captured at 21°C.

ImageJ was used to combine two 500-frame videos of the same region of interest to create the final 1000 frame videos. Videos were analyzed using FIESTA software (30), which fitted two-dimensional Gaussians to the point spread functions of the Tetraspeck beads and the Qdot-labeled cellulases to create single-molecule trajectories. The resulting traces were imported into scripts written in MATLAB (to be submitted to Github) for further analysis of individual tracks. The position changes of Tetraspeck beads were subtracted from all tracks to correct for stage drift in the x-y direction. Particles with total binding durations of less than 10 sec were not included in the analysis since it was often difficult to differentiate processive segments from spatial variances observed in static segments. To avoid premature termination of binding events, 1000 s videos were recorded, but only particles that landed in the first 500 s were analyzed. Few molecules had binding durations greater than 510 s, but those that did were excluded from analysis, as they were potentially due to irreversible binding by denatured enzymes, and thus were considered outliers. For the dwell times of static segments before and after processive movement, only segments with durations more than 5 sec and less than 310 sec were used in the single exponential fit because at least 5 sec were required to determine which state the enzyme was in and few segments had durations longer than 310 sec.

MEMLET was used to fit exponential distributions of dwell times and binding durations by Maximum Likelihood Estimation(39). A single exponential was fit to the data to provide the average value for the various parameters based on a single exponential function applied to the histogram. 95% confidence intervals were obtained by bootstrapping, using 1000 iterations of data selection with replacement.

## Acknowledgements

The authors thank Dannielle Gibson for technical assistance, Scott Pflumm for work in establishing the single-molecule assays, and Haw Yang for support. The authors also thank the Penn State CSL Behring Fermentation Facility - University Park, PA for assistance with cellulose preparation. This work was supported by the Department of Energy Office of Science grant number DE-SC0019065 and DE-SC0019364.

